# Genetic and lipidomic identification of tuberculostearic acid as a controller of mycobacterial membrane compartmentalization

**DOI:** 10.1101/2022.08.17.504266

**Authors:** Malavika Prithviraj, Takehiro Kado, Jacob A. Mayfield, David C. Young, Annie D. Huang, Daisuke Motooka, Shota Nakamura, M. Sloan Siegrist, D. Branch Moody, Yasu S. Morita

**Author notes:** To whom correspondence should be addressed: Department of Microbiology, University of Massachusetts, 639 North Pleasant Street, Amherst MA 01003, USA. These authors contributed equally to this work (Malavika Prithviraj, Takehiro Kado, Jacob A. Mayfield).

## Abstract

Mycobacteria diverge in a basic way from other bacterial and eukaryotic cells based on their distinct membrane structures. Here we report genome-wide transposon sequencing to discover the controllers of membrane compartmentalization in *Mycobacterium smegmatis. cfa*, a gene that encodes a putative cyclopropane-fatty-acyl-phospholipid synthase, shows the most significant effect on recovery from a membrane destabilizer, dibucaine. Lipidomic analysis of *cfa* deletion mutants demonstrates an essential role of Cfa in the synthesis of specific membrane lipids containing a C19:0 monomethyl-branched stearic acid. This molecule, also known as tuberculostearic acid (TBSA), has been intensively studied for decades due to its high level and genus-specific expression in mycobacteria. The proposed Cfa-mediated conversion of an unsaturation to a methylation matched well with its proposed role in lateral membrane organization, so we used new tools to determine the non-redundant effects of Cfa and TBSA in mycobacterial cells. *cfa* expression regulated major classes of membrane lipids including phosphatidylinositols, phosphatidylethanolamines and phosphatidylinositol mannosides. Cfa localized within the intracellular membrane domain (IMD), where it controls both cellular growth and recovery from membrane fluidization by facilitating subpolar localization of the IMD. Overall, *cfa* controls lateral membrane partitioning but does not detectably alter orthogonal transmembrane permeability. More generally, these results support the proposed role of the subpolar IMD as a subcellular site of mycobacterial control of membrane function.

**Significance:** Mycobacteria remain major causes of disease worldwide based in part on their unusual membrane structures, which interface with the host. Here we discover the long sought biosynthetic origin of tuberculostearic acid (TBSA), a major fatty acid found selectively in mycobacteria, as well as its role in mycobacterial cells. The lipid is produced by an enzyme called Cfa, whose loss causes a growth defect and slow reformation of a membrane domain near the pole of the rod-shaped cell. Thus, our study offers mechanistic insights to the intrinsic molecular factors critical for mycobacterial plasma membrane partitioning.

## Introduction

Mycobacteria are encapsulated by a thick and waxy cell envelope, which acts as a barrier to antibiotics and host immunity. Mycobacterial cell envelopes are worthy of investigation because of the high worldwide burden of infectious disease caused by mycobacteria. Also, mycobacterial membranes have evolved structural adaptations that are fundamentally different from eukaryotic cells and model eubacterial organisms based on the number and nature of cell envelope layers present. The mycobacterial cell envelope consists of five biochemically distinct layers: the plasma membrane, the peptidoglycan layer, the arabinogalactan layer, the mycomembrane, and the capsule (1–4). The high impermeability of the mycobacterial cell envelope is attributed to long-chain mycolic acids of the outer mycomembrane. However, the inner plasma membrane may also contribute to the regulation of the permeability of mycobacterial cellular response (5). Indeed, our recent study suggests that mycobacteria have a rapid stress response mechanism to remodel the plasma membrane after exposure to membrane fluidizing chemicals (6). The main building blocks of the plasma membrane are phospholipids; where cardiolipin, phosphatidylethanolamine (PE), phosphatidylinositol (PI), and PI mannosides (PIMs) are the major phospholipid species. Finetuning the composition of phospholipid headgroups and hydrocarbon chains ensures the overall integrity of plasma membrane (7).

Further, recent work makes clear that the mycobacterial plasma membrane is laterally heterogenous and actively regulated. Density gradient fractionation of cell lysate revealed two physically separable fractions with distinct densities and content. One, containing plasma membrane free of cell wall components, is called the intracellular membrane domain (IMD). A second denser fraction, containing <u>p</u>lasma <u>m</u>embrane tightly associated with <u>c</u>ell <u>w</u>all components, is called the PM-CW (8, 9). The proteome and lipidome of the IMD is distinct from those of the PM-CW; the IMD harbors enzymes that are important for active growth and homeostasis, suggesting that the IMD is a biosynthetic hub in the bacteria (8). Consistent with a role in active growth, the IMD is enriched in the subpolar region where rod-shaped mycobacteria grow and elongate (10–12).

In non-growing cells, the IMD is spatially reorganized and delocalized from the subpolar regions to more proximal columnar parts of the cell (13). IMD delocalization was also observed under nutrient starvation, stationary growth, and upon cell wall-targeted antibiotic treatment. Furthermore, membrane-targeted perturbations induced by fluidizers such as benzyl alcohol and dibucaine disrupted the subpolar enrichment of the IMD (14, 15). Thus, emerging data suggest that mycobacteria have a mechanism to spatiotemporally coordinate the IMD in response to stress and different growth conditions.

To gain insight into the molecular mechanism of plasma membrane partitioning in mycobacteria, we screened for genes that are critical for *Mycobacterium smegmatis* to survive and recover from dibucaine treatment. Dibucaine is a topical anesthetic, which preferentially inserts into liquid-ordered phase of a membrane to fluidize the membrane and perturb the lateral membrane organization (16). Here we use unbiased whole organism genetic and lipidomic screens and targeted gene deletion to discover that cyclopropane-fatty-acyl-phospholipid synthase (encoded by *cfa*) controls membrane recovery response and mycobacterial growth *in vitro*. Mechanistic investigation demonstrates that Cfa localizes to the IMD, where it plays an essential role in producing an abundant and mycobacteria-characteristic lipid known as tuberculostearic acid (TBSA) that is distributed among the major membrane phospholipids and controls membrane compartmentalization and growth of *M. smegmatis*.

## Results

### Dibucaine is bacteriostatic and transiently delocalizes IMD from the subpolar regions

The treatment of *M. smegmatis* with 200 μg/mL dibucaine for 3 hours disrupts the polar localization of IMD-associated proteins such as MurG-Dendra2, mCherry-GlfT2, and Ppm1-mNeonGreen (14, 15). Using a previously established strain, in which mCherry-GlfT2 and Ppm1-mNeonGreen are expressed from the endogenous loci (8), we first measured bacterial growth and IMD marker dispersion over time during dibucaine treatment. In a 3-hour window in which we observed a slight, non-significant viability decline (**Fig. S1A**), localization of both mCherry-GlfT2 and Ppm1-mNeonGreen was selectively affected at the poles and progressively diminished over 3 hours (**Fig. 1A**). DivIVA is a pole-associated non-IMD control membrane protein, which was unaffected by dibucaine (**Fig. S1B**). Selective effects on known IMD markers suggest that dibucaine specifically disrupts the IMD.

**Figure 1.**
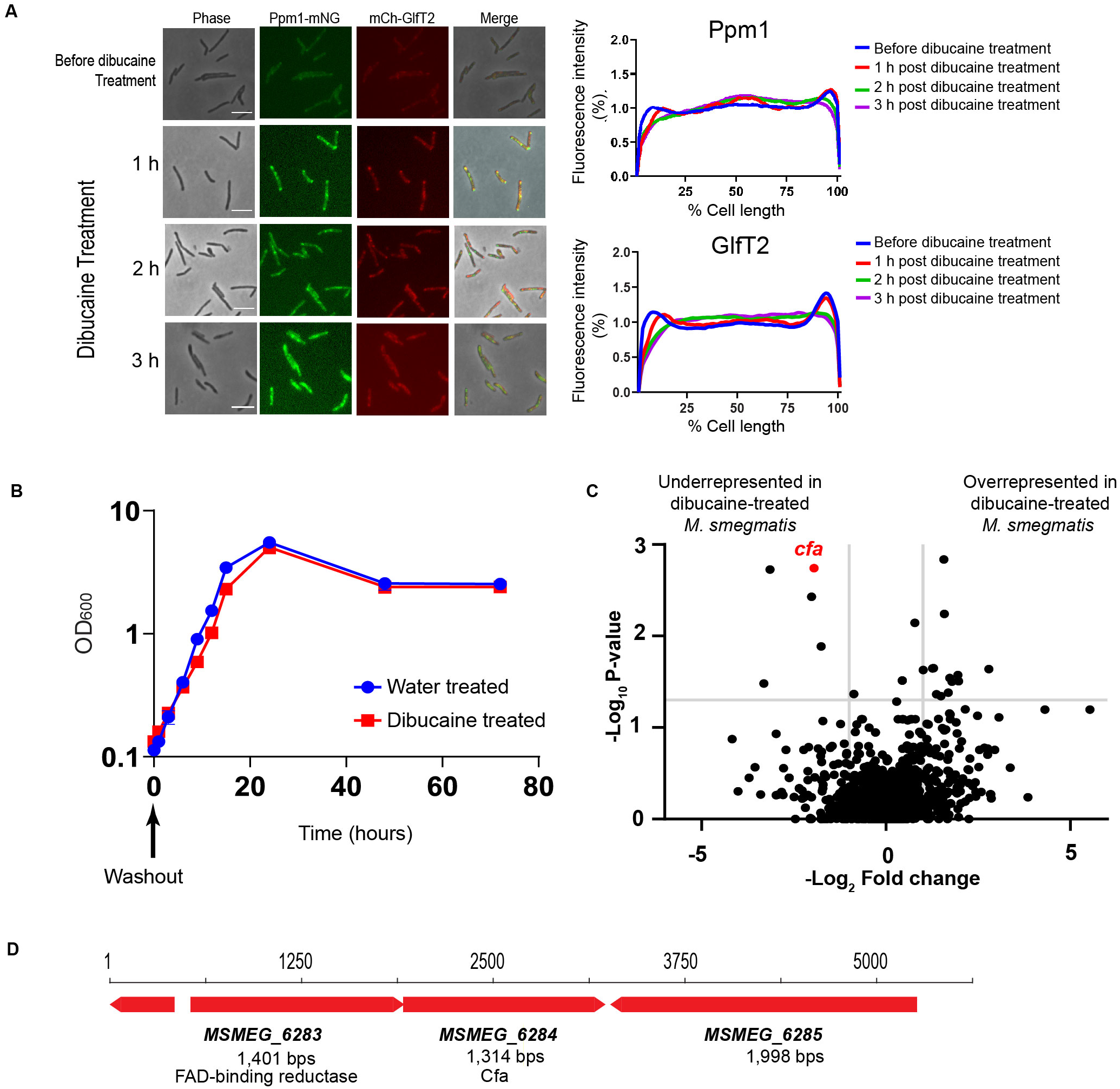
Tn-Seq identifies *cfa* as a gene critical for surviving or recovering from dibucaine treatment. (A) Effect of dibucaine on IMD-associated proteins. IMD-marker proteins, mCherry-GlfT2 and Ppm1-mNeonGreen, were observed every hour during dibucaine treatment (left). Scale bar, 5 μm. Fluorescence intensity profiles along the long axis of the cells were quantified by Oufti (53) (right). Ppm1: n = 237 (before), 121 (1 h), 289 (2 h), and 172 (3 h). GlfT2: n = 127 (before), 119 (1 h), 287 (2 h), or 169 (3 h). (B) Recovery from dibucaine treatment. The same strain expressing mCherry-GlfT2 and Ppm1-mNeonGreen was treated with dibucaine for 3 h, washed, and recovered in a fresh Middlebrook 7H9 medium. The growth recovery was monitored by OD_600_. Time 0 corresponds to the beginning of the recovery period. (C) Transposon mutant library was treated with dibucaine or water, washed, and recovered for ~16 hours. Genes underrepresented or overrepresented in cells treated with dibucaine vs. water are shown in the volcano plot. P-values were calculated by Mann Whitney U-test from 3 independent experiments. The horizontal grey line indicates p = 0.05; the vertical grey lines indicate 2-fold change. (D) The operon structure of *cfa* and the upstream gene encoding FAD-binding reductase in *M. smegmatis*.

Although the subpolar enrichment of IMD was diminished, the IMD was still biochemically detectable after sucrose density gradient fractionation (**Fig. S1C**). However, the distribution of IMD marker proteins, Ppm1-mNeonGreen and PimB’, became more diffuse, again suggesting IMD disruption. MptC, a PM-CW marker, was unaffected. We then used a 3-hour dibucaine pulse followed by wash and chase to examine recovery by growth rate (OD_600_). Dibucaine showed little effect on growth and cells resumed a normal rate of growth almost immediately (**Fig. 1B**). These results show that dibucaine disrupts IMD polarization and attenuates cell replication, but these bacteriostatic effects dissipate when dibucaine is removed.

### Cfa protects against dibucaine stress

To discover the genes mediating resistance to dibucaine treatment, we treated a transposon (Tn) library of the *M. smegmatis* strain expressing mCherry-GlfT2 and Ppm1-mNeonGreen, with dibucaine for 3 hours. We compared Tn insertion frequencies in dibucaine-treated cells with those in vehicle-treated cells. The volcano plot showed five genes significantly diminished in frequency after dibucaine treatment (**Fig. 1C, Table S1**). Among them, *cfa* (*MSMEG_6284*) was depleted approximately 4-fold (P-value, 0.0018) in dibucaine-treated cells in comparison to vehicle-treated cells, making it the most significant genetic change detected and suggesting that *cfa* is important for either surviving during or recovering from dibucaine treatment.

The *cfa* gene encodes a putative cyclopropane fatty acyl phospholipid synthase and forms a putative operon with the upstream *MSMEG_6283* gene annotated as a flavin adenine dinucleotide (FAD)-binding domain protein (**Fig. 1D**). The *cfa* operon structure is widely conserved among mycobacteria, including the key pathogen, *Mycobacterium tuberculosis*, as well as *Mycobacterium leprae*, which is likewise pathogenic and has otherwise undergone massive restriction in its genome size compared to other mycobacteria (17) (**Fig. S2**). In *Escherichia coli*, heterologous expression of this operon from *Mycobacterium chlorophenolicum* resulted in the production of 10-methyl octadecanoic acid, also known as tuberculostearic acid (TBSA), from oleic acid (18). Moreover, one study reported the presence of TBSA in all 61 strains of *M. tuberculosis* complex and 47 strains of nontuberculous mycobacteria except *Mycobacterium gordonae* (19), and this operon is apparently absent in the whole genome of *M. gordonae* (**Fig. S3**). These studies imply the potential role of Cfa in TBSA synthesis, but a role has not been genetically tested in mycobacteria, and any essential function of CFA in mycobacterial cells remains unknown.

### Cfa detectably alters PIMs

To determine the function of *cfa*, we obtained a Δ*cfa* mutant from the Mycobacterial Systems Resource, and complemented it with an expression vector for Cfa-Dendra2-FLAG (Δ*cfa* L5::*cfa-dendra2-flag*, or Δ*cfa* c for short). When grown in a standard rich medium (Middlebrook 7H9) without dibucaine stress, the mutant did not show any significant growth defects (**Fig. S4A**). Cfa was previously identified as an IMD-associated protein by high-throughput proteomic analysis and fluorescence microscopy (8, 20). Indeed, under fluorescence microscopy, Cfa-Dendra2-FLAG was specifically enriched in the subpolar region with sidewall patches (**Fig. S4B**), indicating the IMD association of the protein. In our previous study, Cfa was lost from the IMD proteome when the IMD membrane vesicles were purified by immunoprecipitation (8), potentially implying weak association with the IMD. Consistent with this observation, Cfa-Dendra2-FLAG was identified in both the IMD and cytosolic fractions by density gradient fractionation analysis (**Fig. S4C**).

Next, we extracted lipids and analyzed by high-performance thin layer chromatography (HPTLC). Major membrane phospholipids (**Fig. S4D**), including phosphatidylinositol (PI), phosphatidylethanolamine (PE), cardiolipin (CL), as well as glycopeptidolipids, and trehalose dimycolate (**Fig. S4E**) were equivalent in density after *cfa* knockout. Interestingly, tetraacylated species of PIMs, Ac_2_PIM2 and Ac_2_PIM6, increased after *cfa* knockout (**Fig. S4F**). This outcome of altered PIM pools after gene deletion matches the separately observed response of PIMs to membrane fluidizers (6, 14). Combined, these results, as well as the emergence of *cfa* from a membrane fluidizer-based screen (**Fig. 1C**), provided a hint for a candidate function of Δ*cfa* in control of membrane fluidization in some way that involves membrane lipids.

### Global lipidomics of *cfa* mutants

TBSA is a major fatty acid among *M. smegmatis* phospholipids (21, 22). In particular, PIMs exclusively carry TBSA at *sn*-1 position of glycerol (23). If Cfa is involved in TBSA synthesis, the lack of *cfa* will result in changes in fatty acid composition, which may not be detected by low resolution HPTLC analysis of bulk lipids. We therefore conducted an unbiased high performance liquid chromatography mass spectrometry (HPLC-MS)-based analysis of total lipid extracts from wildtype, Δ*cfa*, and Δ*cfa* c in biological quadruplicate. This lipidomics platform broadly measures named phospholipids, neutral lipids and many hundreds of unnamed lipids that can be tracked across genetically altered samples based on their accurate mass retention time values (**Fig. 2A**). Among 2442 identified ion chromatograms, 366 ions were significantly (corrected p< 0.05, fold-change >2) enriched in Δ*cfa*, while 453 were enriched in wildtype and Δ*cfa c*. Thus, in contrast to the narrow spectrum of PIM change seen in HPTLC, the more sensitive MS-based lipidomics approach demonstrated a broader scope of lipidic change. These many lipid changes represented a potential explanation for the observed effects of *cfa* on recovery from membrane disorder, as well as an opportunity to discover *cfa* as a putative lipid modifying gene.

**Figure 2.**
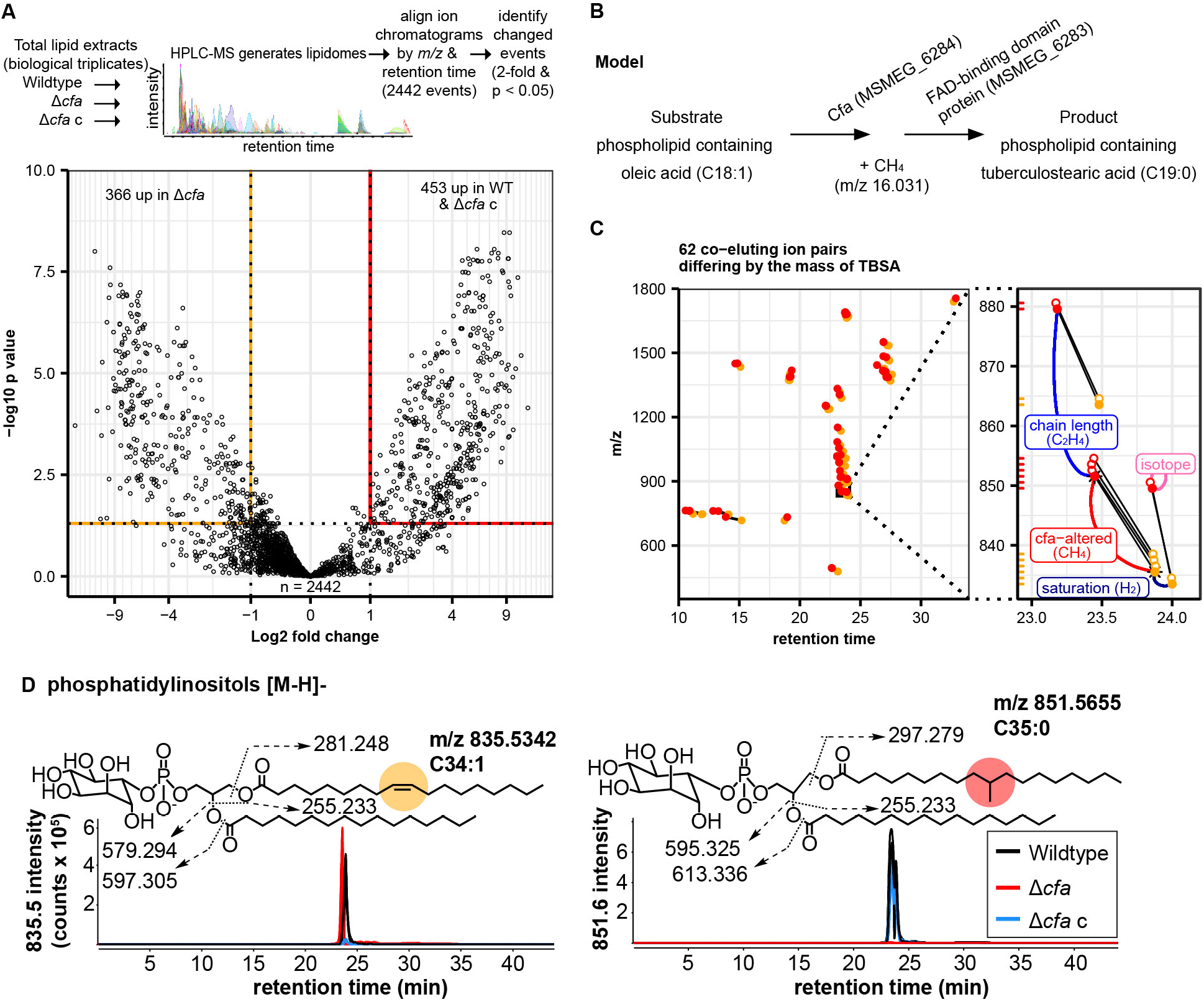
Comparative lipidomes from wildtype *M. smegmatis*, Δ*cfa* and its genetic complement. (A) Schematic of a comparative lipidomic comparison and volcano plot of lipid ions differentially accumulating in Δ*cfa* versus wildtype and Δ*cfa* c detected by negative mode HPLC-MS. Open circles indicate individual ions. (B) Model for Cfa enzymatic function consistent with the observed mass differences. (C) Mass versus retention time for 62 nearly co-eluting lipid pairs identified based on a CH_4_ mass change. Red circles indicate lipids enriched in wildtype and Δ*cfa* c while orange circles indicate lipids enriched in Δ*cfa*. Lipid pairs differing by a CH_4_ mass change and < 1.5 minutes are connected by line segments. Inset, expansion of the plot area indicated by the black rectangle in the main plot showing mass values matching an [M-H]- of C19:0, C16:0 PI (observed *m/z* 851.5653; red closed over asterisk with red label) and its cognate C18:1, C16:0 PI pair (observed *m/z* 835.5333; orange closed over asterisk). Pairs of a chain length variant, an unsaturation variant, and isotopes (open) are labelled. Mass intervals are also shown by a y-axis marginal rug plot. (D) Targeted analysis of lipids regulated by *cfa*. Ion chromatograms of PI containing C18:1, C16:0 (left) or C19:0, C16:0 (right) showing opposite effects of *cfa* deletion on the proposed precursor and product of Cfa. CID-MS interpreted collisional diagrams in each panel show the fragments observed by CID-MS within 10 ppm of the expected exact mass and are diagnostic for identification. Positions of fatty acid attachment, unsaturation (orange circle) and methylation (red circle) are inferred from the literature.

### Comparative lipidomics query for loss of TBSA

To identify the lipids for which *cfa* is essential, we sought targeted analysis of lipids downregulated by *cfa* deletion. First, we formulated a lipidomic query based on the arithmetic difference between the mass of oleic acid (m/z 282.256) and TBSA (m/z 298.287), which are the putative precursors and products of Cfa. If Cfa is essential for TBSA (C19:0) biosynthesis, substitution with oleic acid (C18:1) would reduce the overall mass of any TBSA-containing lipid by CH_4_ (m/z 16.031) (**Fig. 2B**), and the two lipids would nearly co-elute. Indeed, 62 lipid pairs met these criteria in the comparison of Δ*cfa* versus wildtype and Δ*cfa* c (**Fig. 2C**), ranging from m/z 450 to 1800 and in a retention time range of 10 to 33 minutes. This degree of retention is greater than seen for neutral lipids, but matches the general range seen for phospholipids (24). Overall, this pattern suggested broad substitution of C18:1 for C19:0 in many types of polar lipids after deletion of *cfa*.

### Identification of *cfa*-dependent lipids

Next, we sought to identify key *cfa*-dependent lipids from these ion pairs. The *M. smegmatis* lipidome is not yet annotated, but we could use mass-based annotations for lipids shared with the *M. tuberculosis* lipidome (24, 25) and confirm chemical structures with collision-induced dissociation mass spectrometry (CID-MS). Given the large number of lipids affected by *cfa* deletion, we simplified the analysis by taking advantage of the phenomenon whereby structurally similar lipids cluster into groups with similar mass and retention time (**Fig. 2B-C**). Then, we implemented a strategy to chemically solve one compound in each cluster based on embedded mass values and then deduce the structures corresponding to all ions in one group. For example, one member of a *cfa*-dependent ion pair matched the expected mass (m/z 851.566) of PI comprised of TBSA and palmitic acid (PI C19:0, C16:0). This lipid clustered together with a PI compound matching the mass of a chain length variant (PI C19:0, C18:0), a saturation variant (PI C19:0, C16:1) and 6 isotopes of these molecules (**Fig. 2C, inset**). We ruled in the PI structure with CID-MS that detected glyceryl-inositol-phosphate and the defining fatty acyl fragments (**Fig. 2D**). The strong signal of PI C19:0, C16:0 in ion chromatograms (~600,000 counts) was consistent with high-level biosynthesis (**Fig. 2D**). Ion chromatograms also demonstrated the complete loss of PI C19:0, C16:0 in Δ*cfa* and its restoration through complementation, establishing the essential role of *cfa* in its biosynthesis. Importantly, while *cfa* was essential for the TBSA-containing form of PI, deletion increased the production of PI formed from C18:1 oleic acid, which is the putative precursor of TBSA (**Fig. 2D**). All outcomes could be explained if *cfa* encodes the essential enzyme for converting C18:1 oleic acid to C19:0 TBSA.

Extending this analysis of putative *cfa*-dependent ion pairs to other major membrane phospholipids, *cfa*-regulated lipid pairs were identified as PE *(m/z* 716.5326) and AcPIM2 (*m/z* 1397.8695) by CID-MS (**Fig. S5A-B**). Similar to PI species, the *cfa*-dependence of ion chromatograms of PE and AcPIM2 variants containing C19:0 fatty acids was complete and genetically complementable, demonstrating an essential role for *cfa*. Also, both PE and AcPIM2 showed strong increases in forms containing C18:1 fatty acids, which ruled out the possibility of a general block of PE and PIM biosyntheses and instead localized the defect to the biosynthesis of TBSA-containing lipids. In summary, the deletion of *cfa* is associated with a selective defect in TBSA incorporation into many mycobacterial lipid families (**Fig. 2 and S5**), including at least three major membrane phospholipids, while leaving the larger total pools of these phospholipids containing other fatty acids intact (**Fig. S4**).

### Effects of *cfa* on mycobacterial growth and membrane integrity

To determine the physiological function of Cfa in live cells, we next tested if the mutant is more susceptible to dibucaine treatment. We treated Δ*cfa* with 200 μg/ml dibucaine for 3 hours and examined the viability of the mutant. Δ*cfa* did not show any increased sensitivity to dibucaine and CFU did not decline from pre-to post-treatment (**Fig. 3A**). We next examined the recovery after dibucaine treatment in a pulse-chase experiment and found that the growth rate of Δ*cfa* was slower than that of the wildtype initially, but the mutant recovered during culture in dibucaine-free media after 15-18 hours (**Fig. 3B**). Thus, Δ*cfa* is defective in recovering from, but not in surviving during, the dibucaine treatment under the conditions tested. We tested another membrane fluidizer, benzyl alcohol, and found that Δ*cfa* was not defective in recovering from this membrane fluidizer (**Fig. 3C**), suggesting that the effect of dibucaine on mycobacterial plasma membrane is distinct from that of benzyl alcohol.

**Figure 3.**
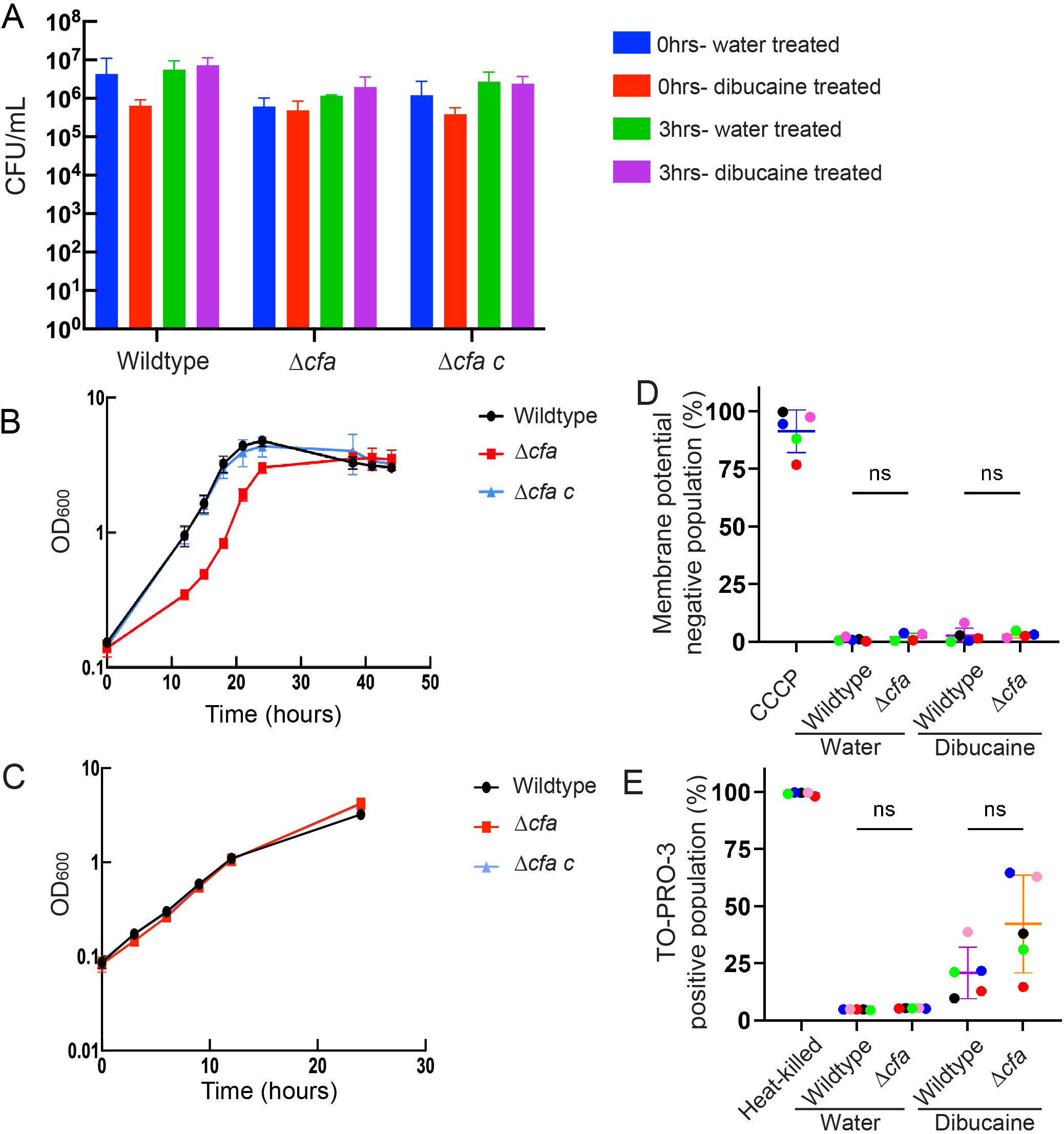
Δ*cfa* shows delayed recovery after dibucaine treatment, but membrane permeability is maintained. (A) CFUs were calculated in biological triplicate to determine the survival of Δ*cfa* during dibucaine treatment. (B and C) Growth after washing out dibucaine (B) or benzyl alcohol (C) was monitored by OD_600_. (D and E) A population of wildtype or Δ*cfa* cells that were negative for membrane potential (measured by the fluorescence ratio (red/green) of DiOC2(3)) (D) or positive for membrane permeability (measured by TO-PRO-3 stains) (E) was plotted. Carbonyl cyanide chlorophenylhydrazone (CCCP) was used to disrupt membrane potential. Heat (65°C, 1 h) was used to permeabilize membrane. Each color shows a set of data from a single replicate. Statistical significances were determined by the Kruskal–Wallis test, followed by Dunn’s multiple comparison test. ns, not statistically significant.

Depletion of TBSA and accumulation of cis-monounsaturated fatty acids in membrane phospholipids predict increased membrane fluidity, which could explain the slower recovery from membrane fluidizing stress. At the extreme, excess membrane fluidity can compromise the permeability barrier such that compounds can orthogonally transit through the plasma membrane (26, 27). To test if the plasma membrane of Δ*cfa* is more permeable, we used the membrane-impermeable DNA staining dye TOPRO3 and the membrane potential probe 3,3′-diethyloxacarbocyanine iodide (DiOC2(3)). As a positive control, heat-killed cells became TOPRO3-positive and lost membrane potential (**Fig. S6**). In contrast, cells lost membrane potential upon treatment with carbonyl cyanide m-chlorophenyl hydrazone (CCCP), a membrane potential uncoupler, while they remained TOPRO3-negative (**Fig. S6**). We compared the wildtype and Δ*cfa* with or without dibucaine treatment in five experiments, which reproducibly found that both wildtype and Δ*cfa* maintained the membrane potential even after dibucaine treatment (**Fig. 3D and S6**). While subpopulations (10-60%) of both wildtype and mutant cells became TOPRO3-positive after dibucaine treatment (**Fig. 3E and S6**), there was no statistically significant difference between the strains (**Fig. 3E**). Taken together, Δ*cfa* is defective in recovering growth after membrane fluidizing treatments, but plasma membrane integrity is not substantially compromised to allow orthogonal transit of measured molecules.

### Membrane domain partitioning is altered by *cfa* deletion

Our results so far showed that, in the absence of dibucaine challenge, Δ*cfa* is not overtly defective in growth or its plasma membrane integrity. Since Δ*cfa* was initially identified as a mutant sensitive to dibucaine, which alters membrane compartmentalization, we wondered if Δ*cfa* is defective in recovering from dibucaine-induced disruption of membrane partitioning. Biochemically, both the IMD and PM-CW were purified from Δ*cfa* under normal growth conditions (**Fig. 4A**). However, after 3-hour dibucaine treatment, the IMD marker PimB’ became diffuse and less enriched in the IMD fractions (**Fig. 4A**) like wildtype (see **Fig. S1C**). We then examined the recovery of the polar IMD enrichment over time after dibucaine treatment. In wildtype cells, polar enrichment of the IMD marker mCherry-GlfT2 was partially restored after a 3-hour recovery, and fully restored after 6 hours (**Fig. 4B and 4C**). In contrast, for Δ*cfa*, the polar enrichment of mCherry-GlfT2 required 12 hours for full recovery (**Fig. 4B and 4C**). Thus, Δ*cfa* is delayed in recreating subpolar membrane partitioning after dibucaine treatment. This result is mechanistically notable because the experimentally induced membrane fluidization has more significant impact on the lateral membrane compartmentalization of the mutant when its plasma membrane is predicted to be more fluid due to the substitution of TBSA-containing lipids with lipids containing cismonounsaturated fatty acids.

**Figure 4.**
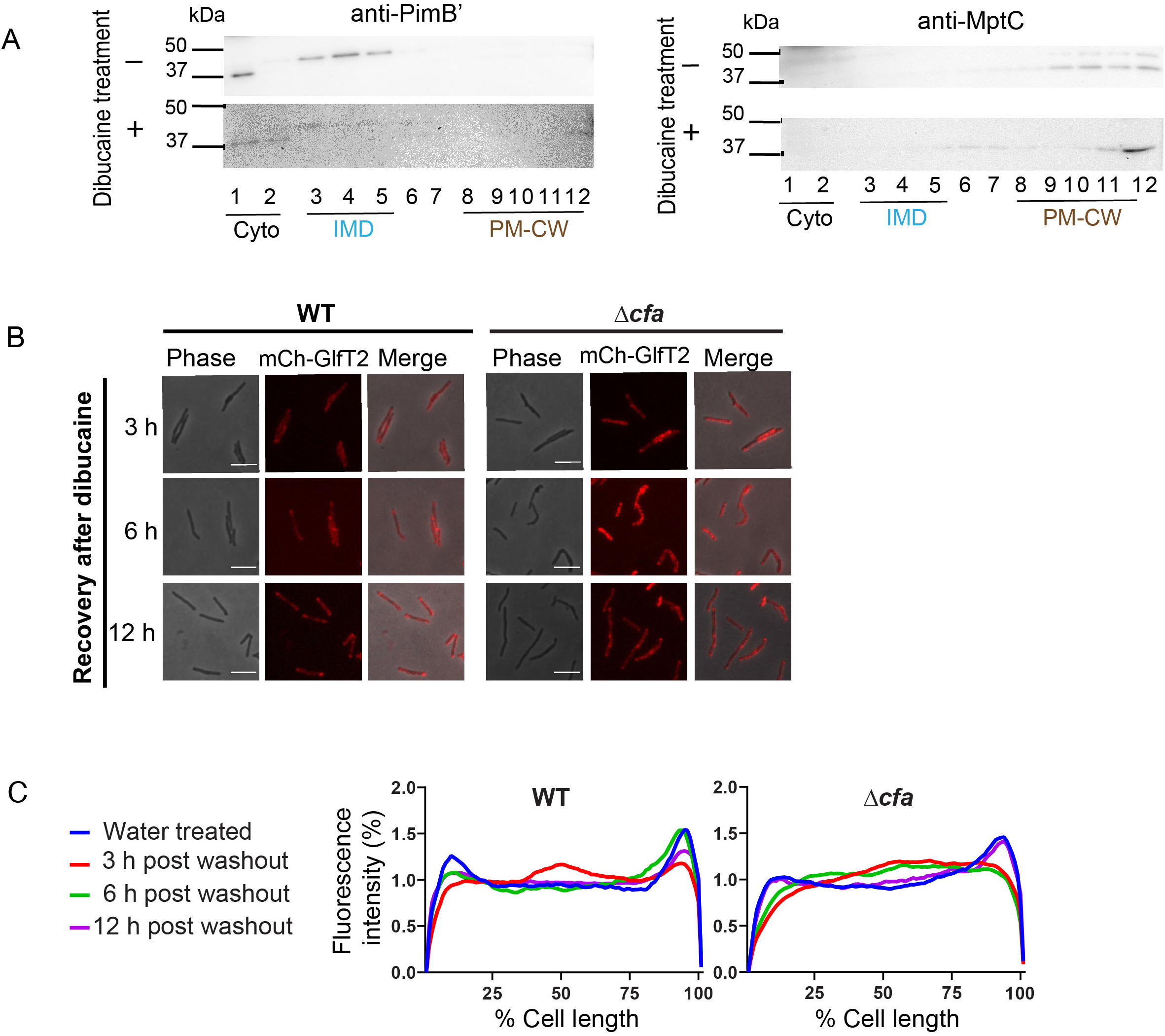
Δ*cfa* is defective in restoring subpolar membrane partitioning. (A) Western blotting of density gradient fractions of Δ*cfa* cell lysates. Before or after 3-hour dibucaine treatment, log phase cells were lysed and fractionated by sucrose density gradient sedimentation. PimB’, an IMD marker; MptC, a PM-CW marker. (B) Recovery of the subpolar localization of the IMD marker mCherry-GlfT2 (mCh-GlfT2) expressed from the endogenous locus in wildtype (WT) or Δ*cfa* cells. Cells were treated with 200 μg/ml dibucaine for 3 hour, washed, and recovered for up to 12 hour. Images were taken at 3, 6, and 12 hour. Scale bar, 5 μm. (C) Profiles of relative fluorescence intensities from images taken as in panel B. Wildtype: n = 46 (water), 93 (3 hour), 54 (6 hour), or 92 (12 hour). Δ*cfa*: n = 75 (water), 205 (3 hour), 138 (6 hour), or 77 (12 hour).

## Discussion

Tuberculostearic acid (TBSA) has been studied for decades as an abundant mycobacteria-characteristic lipid. TBSA is produced by most mycobacterial species, where it comprises from 10 to 20 percent of fatty acids (28, 29). Because TBSA is not produced by humans, it has been proposed as a diagnostic test for tuberculosis (30–36). The structure of TBSA was determined in 1934 (37), and it was first synthesized in 1947 (38). TBSA biosynthesis from oleic acid was proposed in 1962 (39), which frames the interpretations of the cellular lipidomics performed here.

The operon structure of *cfa* and the upstream gene encoding an FAD-binding oxidoreductase is well conserved among mycobacteria. Furthermore, heterologous expression of the two-gene operon from *M. chlorophenolicum* in *E. coli* resulted in an accumulation of lipids carrying a TBSA in *E. coli* (18). However, direct demonstration that Cfa is for TBSA synthesis in mycobacteria was lacking and the precise identity and number of genes with this function was unknown. There are seven and nine paralogs of *cfa* in *M. smegmatis* and *M. tuberculosis*,respectively, and at least two other genes have also been proposed to mediate the biosynthesis of TBSA in mycobacteria (40, 41). Our study shows that *cfa* (*MSMEG_6284* in *M. smegmatis*), rather than the other two previously proposed genes, is the essential gene needed for the biosynthesis of many TBSA-containing lipids, including three major families of membrane phospholipids PI, PE and PIMs (23, 30, 42). The defects of Δ*cfa* that is restored with *cfa* complementation unequivocally demonstrates the necessary and sufficient role of Cfa in TBSA synthesis. Given the prior work by Machida *et. al*. (18), we speculate that the FAD-binding oxidoreductase likely mediates the second step of the reaction, where cis-9,10-methylene octadecanoic acid produced by Cfa is reduced to TBSA.

Through comparative lipidomics, unbiased examination of all ionizable lipids altered through gene deletion in live mycobacterial cells is now possible (43), generating a dominant pattern of CH4 loss among polar lipids. Mechanistically, we strongly favor the interpretation that *cfa* deletion causes loss of these lipids through ablation of the shared TBSA pool, rather than some adjunct effect of Cfa on overall PI, PE, and PIM synthesis. Lipidomics data demonstrated both the ablation of TBSA-containing phospholipids and increased C18:1-containing PE, PI, and AcPIM2. The opposite effects of *cfa* deletion based on the fatty acid present ruled out a general block in synthesis of these phospholipid products, and it strongly supports that *cfa* deletion ablated the downstream product (TBSA) and increased the immediate upstream product (oleic acid). Also, the opposite regulation of distinct PI and PE acyl forms by *cfa* can resolve the otherwise apparent contradiction between preserved total PI and PE pools observed in HPTLC and the complete loss of TBSA-containing PI and PE in MS-based lipidomics.

In *Mycobacterium phlei*, the fatty acid composition shifts from TBSA to cis-monounsaturated fatty acids such as oleic and linoleic acids when grown at a lower temperature (29). This observation implies that tilting the balance to cis-unsaturated fatty acid from TBSA helps the cells to maintain membrane fluidity under a colder growth temperature, where the membrane becomes more ordered. Therefore, TBSA is likely involved in homeoviscous adaptation, a stress response mechanism (44, 45), in which TBSA makes the plasma membrane more rigid than cis-unsaturated fatty acids. Our data are consistent with the importance of TBSA in homeoviscous adaptation as the lack of TBSA and the accumulation of oleic acids in Δ*cfa* would make the mutant’s plasma membrane more disordered, explaining the reason why Δ*cfa* is more vulnerable to dibucaine, a membrane fluidizing molecule. We propose that TBSA biosynthesis is important for creating a lipid environment resilient to external threats that increase the fluidity of the mycobacterial plasma membrane.

Branched-chain fatty acids are important for membrane homeostasis and their depletion makes bacterial cells defective in membrane fission and fluidity maintenance (46–48). While these studies used iso and anteiso fatty acids in other bacteria, we propose TBSA in mycobacteria could similarly support membrane integrity. However, our data indicate that the defect in synthesizing TBSA has a specific effect on the kinetics of membrane domain partitioning and not on other functions. First, Δ*cfa* grows like the wildtype, indicating that there are no gross impacts on growth under a laboratory culture condition. Second, we used the membrane potential probe DiOC2(3) and demonstrated that Δ*cfa* can create a proton gradient across the plasma membrane for oxidative phosphorylation. Third, we tested membrane permeability of Δ*cfa* using the fluorescent DNA-binding dye TO-PRO3 and did not observe any significant defects relative to the wildtype with or without dibucaine challenge. These observations suggested that the lack of Cfa and resultant changes in the ratio of oleic acid to TBSA do not have significant effects on overall membrane integrity. In contrast, we found that there was a delay in polar IMD enrichment in Δ*cfa* during recovery from dibucaine treatment. This delay coincided with delayed recovery of growth after dibucaine treatment. Benzyl alcohol has a similar disruptive effect on membrane partitioning (15), and yet the recovery of growth of Δ*cfa* after the treatment with this membrane fluidizer was no different from that of the wildtype. Dibucaine has been suggested to insert into ordered membrane domain preferentially while benzyl alcohol preferentially inserts into disordered regions (16, 49). Thus, we speculate that the molecular mechanisms of disrupting mycobacterial membrane partitioning are different between these two membrane fluidizers. Although TBSA-containing phospholipids are distributed in both the IMD and the PM-CW (8), they could have more dominant roles in maintaining the integrity of the PM-CW, which, we speculate, is more ordered than the IMD (50).

We recently reported inositol acylation of PIMs as a response to membrane fluidization conserved in *M. tuberculosis* (6). Our current study indicates the role of TBSA-containing lipids, which are widely conserved in mycobacteria including *M. tuberculosis*, in homeoviscous adaptation. One transposon mutagenesis study predicts *cfa* to be essential in *M. tuberculosis* (51). Furthermore, the *cfa* ortholog in *Mycobacterium bovis* BCG is critical for persistence during the passage in bovine lymph node (52). These studies illuminate the potential importance of homeoviscous adaptation in *M. tuberculosis*, an obligate human pathogen that does not experience temperature fluctuations. Thus, our studies pinpoint the significance of yet-to-be-defined host factors that affect the pathogen’s membrane fluidity during host-pathogen interactions.

## Materials and Methods

Full descriptions of methods are given in SI Appendix.

## Acknowledgements

This work was supported by NIH to YSM and MSS (R21 AI144748) and to DBM (R01 AI165573 and U19 AI162584). MP was supported by a fellowship from the University of Massachusetts Amherst as part of the Chemistry-Biology Interface Training Program (National Research Service Award T32 GM008515 and GM139789). TK was supported by an Uehara Memorial Foundation Postdoctoral Fellowship. We thank Drs. Eric Rubin and Chidi Akusobi for help with Tn-seq experiments. We thank the support of the Flow Cytometry and Genomics Resource Cores (Directors Drs. Amy Burnside and Ravi Ranjan) at the University of Massachusetts Institute for Applied Life Sciences.

