## Supplemental Information for "Genetic and lipidomic identification of tuberculostearic acid as a controller of mycobacterial membrane compartmentalization"

**Supporting Information Appendix**

**Materials and Methods**

Growth and treatment with membrane fluidizers

*M. smegmatis* mc$^2$155 was grown at 37°C in Middlebrook 7H9 broth supplemented with 11 mM glucose, 14.5 mM NaCl, and 0.05% (v/v) Tween-80. Dibucaine (Millipore-Sigma) or water (as the vehicle control) was added to a log-phase culture at the final concentration of 200 µg/ml. After 3-hour incubation at 37°C, the culture was washed with phosphate-buffered saline (PBS) containing 0.05% Tween-80 (PBST) three times and resuspended in Middlebrook 7H9 broth for recovery. Benzyl alcohol treatment was identical to dibucaine treatment except that the treatment was for 1 hour at the final concentration of 100 mM. Colony forming units were determined by serially diluting cell culture using Middlebrook 7H9 broth and spotting 5 µl on Middlebrook 7H10 agar supplemented with 11 mM glucose and 14.5 mM NaCl. The agar plates were incubated at 37°C for 2-3 days before counting the number of microcolonies.

Transposon sequencing

We prepared a transposon mutant library using ΦMycoMarT7 phage as before (1), grow the library of cells to a log phase, and treated the cells with dibucaine (or water as the vehicle control) for 3 hours. The cells were washed to remove dibucaine and recovered in Middlebrook 7H9 broth for ~16 hours. Genomic DNA was purified, sheared, and barcoded. Transposon insertion sites were then amplified by a nested PCR. To prepare the library for high-throughput DNA sequencing, we used the KAPA Library Preparation Kit (Kapa Biosystems) and TruSeq adapters (Illumina) as before (2). The library was sequenced by 100-bp paired-end sequencing using the Illumina HiSeq3000 platform. Identified genes were compared between water- or dibucaine-treated samples using TRANSIT as in the previous literatures (3, 4). Library sequencing yielded 5 million unique transposon-inserted-sequences, which covered over 35% of the possible TA sites in the genome.

Bioinformatic analysis of *M. gordonae* genome region

The genome region of *M. gordonae* strain X7091 chromosome (GenBank sequence ID: CP070973.1; region 6782643 – 6796036) and that of *M. smegmatis* mc$^2$155 chromosome (GneBank sequence ID: CP000480.1; region 6340496 – 6353816) were aligned using NCBI Nucleotide BLAST.

Live cell imaging

Live cells were placed on 1% (w/v in water) agar pad and fluorescent protein localization was visualized using a Nikon Eclipse E600 microscope (100x objective, N.A. 1.30) equipped with an ORCA-ER cooled charge-coupled-device camera (Hamamatsu) and Openlab software 5.5.2 (Improvision). All fluorescence images were taken at the exposure of 4 s with a gain of 4. Fluorescence intensity profiles were quantified as before (1). Briefly, cell shape was contoured using Oufti (5), and each cell was divided to 100 sections along the long axis. Average relative fluorescence intensity was calculated using MATLAB with published scripts (6) and plotted along the normalized cell length.

Preparation of mutants and plasmids

*M. smegmatis* expressing both mCherry-GlfT2 and Ppm1-mNeonGreen from their respective endogenous loci was previously established (7). To visualize DivIVA-eGFP, we used the same *M. smegmatis* strain expressing DivIVA-eGFP and mCherry-GlfT2 as before (8). *M. smegmatis* Δ*cfa* and a vector to express Cfa-Dendra2-FLAG were obtained from MSRdb (9). To express mCherry-GlfT2 in Δ*cfa*, pMUM053 was electroporated to replace the endogenous *glfT2* gene with an engineered gene to express mCherry-GlfT2 as before (7).

Density gradient fractionation, SDS-PAGE and immunoblotting

Subcellular fractionation was performed as before (7, 10). Briefly, cells were grown to a log phase ($OD_{600}$ = 0.5~1.0), harvested, and lysed by nitrogen cavitation. The cell lysate was placed on top of a sucrose gradient (20-50%, w/v) and fractionated by sedimentation at 35,000 rpm on SW-40Ti rotor (Beckman-Coulter) at 4°C. SDS-PAGE and immunoblotting of gradient fractions were as previously described (7, 11).

Lipid extraction, HPTLC, and lipidomics

In quadruplicates, each strain was harvested at a log phase ($OD_{600}$ = 0.5~1.0), and the wet cell pellets were sequentially treated with 20 volumes of chloroform/methanol (2:1, v/v), 10 volumes of chloroform/methanol (1:1, v/v) and 10 volumes of chloroform/methanol (1:2, v/v). Ten percent volume of the combined organic phase was set aside for HPTLC analysis and the remainder was subjected to lipidomic analysis. For HPTLC analysis, combined organic phase was dried under a stream of nitrogen gas, and the dried lipids were resuspended in 5 volumes of chloroform/methanol (1:1, v/v) in reference to the original pellet weight. Ten μl of lipid extracts were spotted onto an HPTLC silica gel 60 sheet (Merck) and chromatographed using chloroform/methanol/13 M ammonia/1 M ammonium acetate/water (180:140:9:9:23, v/v/v/v/v) as a solvent. PIMs were visualized by spraying the HPTLC plate with orcinol-$H_2SO_4$ and baking at 120°C. Phospholipids were visualized using a molybdenum blue spray reagent (Sigma-Aldrich). For the purification of GPLs and TDM, dried lipids were further purified by chloroform/methanol/water (8:4:3, v/v/v) phase partitioning. The organic phase containing GPLs and TDM was dried under a nitrogen gas stream and the dried lipids were resuspended in chloroform/methanol/water (9:1:0.1, v/v/v). Lipids were separated on an HPTLC plate using a solvent containing chloroform/methanol/water (9:1:0.1, v/v/v). GPLs were visualized using an orcinol-$H_2SO_4$. Lipidomic analysis and CID-MS was performed as previously described (7) using an Agilent 1260 Infinity LC system and 6546 QTOF mass spectrometer. Dried lipids were dissolved in 70:30 (v/v) hexanes:isopropanol at 1 mg/ml by dried weight and were separated using a normal-phase Inerstil Diol column (GL Sciences, Tokyo, Japan) with 0.1% formic acid and 0.05% aqueous ammonia added to all solvents. HPLC/MS data were analyzed using MassHunter (Agilent) and R for statistical analysis and data visualizations. Raw data and R code are available upon request.

<u>Flow cytometry</u>

A 2-ml aliquot of log phase cells was treated with 200 µg/ml dibucaine for 3 hours at 37°C with shaking. Cells were then treated with 100 nM TO-PRO-3 (ThermoFisher Scientific) and 30 µM DiOC2(3) (TCI Chemicals). After a 15-min incubation, cells were centrifuged at 2,000x *g* for 5 min and the pellet was resuspended in 2% formaldehyde in PBS for fixation. Fixed cells were washed again and resuspended in PBS for flow cytometry analyses using a three-laser (405 nm, 488 nm, and 640 nm) LSR Dual Special Order Research Product (BD Biosciences). As in a previous report (12), DiOC2(3) was excited using 488 nm laser with red emission detected through one filter set (a long pass (LP) filter 600 nm and band pass (BP) filter 610 nm / 20 nm), and green emission detected through another filter set (LP filter 505 nm and BP filter 525 nm / 50 nm). TO-PRO-3 was excited using the 640 nm laser and detected through a BP filter 670 nm / 30 nm. As a positive control for membrane permeability (TO-PRO-3 labeling), cells were heat-killed for one hour at 65°C. As a positive control for the disruption of membrane potential (DiOC2(3) labeling), cells were incubated with 25 µM carbonyl cyanide m-chlorophenyl hydrazone (CCCP, Sigma Aldrich) for an hour. Data were analyzed using FlowJo 10.0 (FlowJo LLC. BD). The red/green ratio of DiOC2(3) was determined using the Derived Parameter function comparing the DiOC2(3) red and green median fluorescence parameters.

**Supporting Tables**

Table S1. Genes identified by Tn-seq

| Underrepresented in dibucaine-treated *M. smegmatis* | | |
|---|---|---|
| Gene locus | Gene name | Gene description |
| MSMEG_5054 | *lpqG* | Probable lipoprotein |
| MSMEG_5072 | *sigE* | RNA polymerase sigma-70 factor |
| MSMEG_5488c | *mprA* | DNA-binding response regulator |
| MSMEG_6284 | *cfa* | cyclopropane-fatty-acyl-phospholipid synthase |
| MSMEG_6398 | - | antigen 85-A |
| Overrepresented in dibucaine-treated *M. smegmatis* | | |
| MSMEG_0840c | - | Hypothetical protein |
| MSMEG_1567 | - | Hypothetical protein |
| MSMEG_1886 | *desA3* | NADPH-dependent stearoyl-CoA 9-desaturase |
| MSMEG_2772 | - | Amino acid permease |
| MSMEG_3119 | - | Unknown transporter |
| MSMEG_3227 | *pyk* | Pyruvate kinase |
| MSMEG_4192c | - | Hypothetical protein |
| MSMEG_4323 | - | Pyruvate dehydrogenase E1 component |
| MSMEG_5121c | - | Aminotransferase |
| MSMEG_5694 | - | Hypothetical protein |
| MSMEG_6761 | - | Glycerol-3-phosphate dehydrogenase |

**Supporting Figure Legends**

**Figure S1. Effects of dibucaine treatment on *M. smegmatis*.** (A) Colony forming units (CFUs) were calculated before and after a dibucaine treatment of the strain, which expresses mCherry-GlfT2 and Ppm1-mNeonGreen from the endogenous loci. CFUs were obtained from biological triplicate data. (B) DivIVA, a PM-CW marker protein, was observed before and after dibucaine treatment. n = 36 (before) or 37 (3 h). (C) Sucrose gradient fractionation of cell lysates of the strain expressing mCherry-GlfT2 and Ppm1-mNeonGreen. The strain was treated with or without dibucaine. Ppm1-mNeonGreen was visualized by in-gel fluorescence after SDS-PAGE. PimB' and MptC were visualized by western blotting using rabbit anti-PimB' and anti-MptC antibodies. MptC is a PM-CW marker (13), which was unaffected by dibucaine treatment.

**Figure S2. Alignment of Cfa proteins.** Cfa proteins from *M. smegmatis* mc$^2$155, *M. marinum*, *M. chlorophenolicum*, *M. abscessus*, *M. tuberculosis* H37Rv, *M. avium*, and *M. leprae* were aligned using UniProt Align (14).

**Figure S3. Synteny of the genome regions from *M. smegmatis* and *M. gordonae*, showing the absence of the *cfa*-containing operon in *M. gordonae*.** (A) Map of the genome region surrounding the *cfa* gene, spanning 6340496 – 6353816 in *M. smegmatis* (Sequence ID: CP000480.1). (B) Pairwise alignment of the region 6340496 – 6353816 of *M. smegmatis* genome and the region 6782643 – 6796036 of *M. gordonae* genome (Sequence ID: CP070973.1). The genome regions of *M. smegmatis* shaded in blue and brown correspond to MSMEG_6278 and MSMEG_6283-6284, respectively. These genes were missing from the syntenic region of the *M. gordonae* genome. (C) When Cfa (MSMEG_6284) was used for Protein BLAST against the *M. gordonae* genome, the closest homolog showed only 32% amino acid identity. This homolog was the ortholog of MSMEG_1203 (60% amino acid identity), an unrelated protein involved in mycolic acid modification, indicating that the ortholog of *cfa* is absent in the *M. gordonae* genome.

**Figure S4. Lipid composition of Δ*cfa* is comparable to wildtype.** (A) Growth curve for each strain grown at 37°C in Middlebrook 7H9 medium, measured by $OD_{600}$. WT, wildtype. (B) Fluorescence microscopy of the dual marker strain, expressing Cfa-Dendra2-FLAG from the *attB* site of mycobacteriophage L5 and the IMD marker HA-mCherry-GlfT2 from the endogenous locus. Scale bar, 5 µm. (C) Density gradient fractionation of a cell lysate of a dual marker strain. Cyto, cytoplasm. (D) HPTLC analysis of phospholipids visualized by molybdenum blue staining. CL, cardiolipin; PE, phosphatidylethanolamine; and PI, phosphatidylinositol. (E) HPTLC analysis of glycopeptidolipids (GPLs) and trehalose dimycolate (TDM) visualized by orcinol staining. (F) HPTLC analysis of PIMs visualized by orcinol staining.

**Figure S5. Targeted analysis of lipids regulated by *cfa*.** Signals corresponding to C18:1-containing PE (*m/z* 716.5236, panel A) and AcPIM2 (*m/z* 1397.8695, panel B) compared to C19:0-containing species after *cfa* deletion show opposing effects. CID-MS interpreted collisional diagrams in each panel show the fragments observed by CID-MS within 10 ppm of the expected exact mass and are diagnostic for identification. Positions of fatty acid attachment, unsaturation (orange circle) and methylation (red circle) are inferred from the literature.

**Figure S6. Membrane permeability barrier and protein gradient formation are largely unaffected in Δ*cfa*.** Wildtype or Δ*cfa* bacteria were stained with a membrane potential sensor (DiOC2(3)) and a membrane permeability sensor (TO-PRO-3) with or without dibucaine. Membrane potential was assessed by the fluorescence ratio (red/green) of DiOC2(3). Carbonyl cyanide chlorophenylhydrazone (CCCP) was used to disrupt membrane potential. Heat (65°C, 1 h) was used to permeabilize the membrane. The data set is a representative result from 5 experiments.

Figure S1

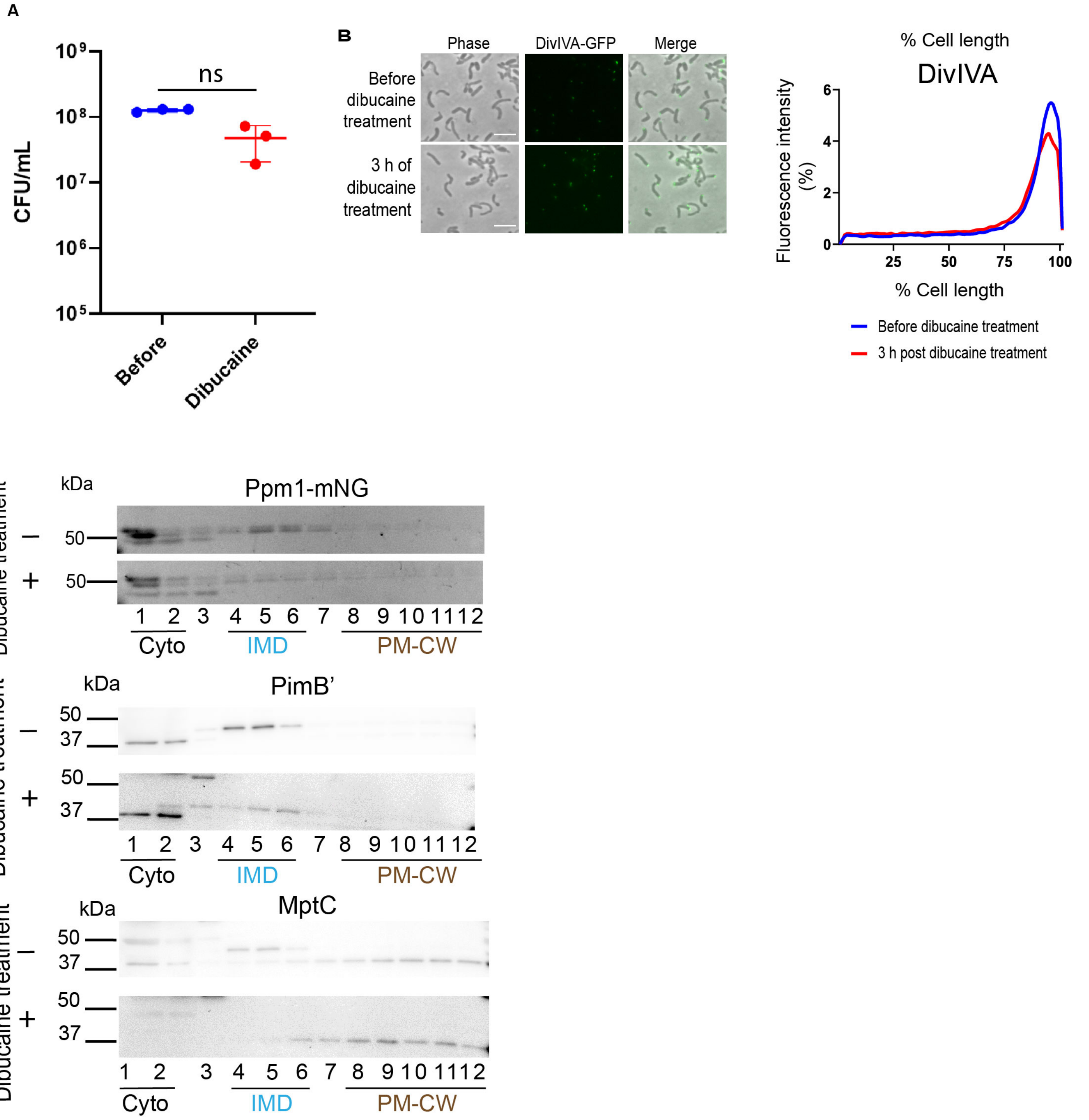
A
ns
CFU/mL
Before
Dibucaine
B
Phase
DivIVA-GFP
Merge
Before dibucaine treatment
3 h of dibucaine treatment
% Cell length
DivIVA
Fluorescence intensity (%)
% Cell length
Before dibucaine treatment
3 h post dibucaine treatment
C
kDa
Ppm1-mNG
Dibucaine treatment
Cyto
IMD
PM-CW
PimB’
MptC

Figure S2

A

```
M. smegmatis        WP_003897704.1     1  MTTFKERETS---TADRKLTLAEILEIFA-AGKEPLKFTAYDGSSAGPEDATMGLDLKTP  56
M. marinum          WP_208024746.1     1  MTATKEA-NQPPNANGGRLSIAEVLAIFXATGXQPLKFSAYDGSTAGCNDAEMGLELTSP  59
M. chlorophinolicum WP_048472121.1     1  MTTFRERPTDAANPADGRLTLAEILEIFA-SGTRPLKFTAYDGSSAGPDDAALGLDLLTP  59
M. abscessus        WP_005065329.1     1  MTTT---RAKNARGENGKLSLAEILELMA-AGDLPLRFSAYDGSTAGPENAELGLDLLTP  56
M. tuberculosis     WP_003899652.1     1  -------------MTTGRLSMAEILEIFTATGQHPLKFTAYDGSTAGQDDATLGLDLRTP  47
M. avium            WP_225520609.1     1  MTTTKQ----PHRSRTGKLSMAEILEVFAATGRHPLKFTAYDGSTAGSEDAELGLDLRSP  56
M. leprae           WP_041323340.1     1  MTTTKDS-ERFTQISSGKLTMAEILGILTATGQQPLKFTAYDGSTAGHDDAEIGLDLCSP  59
                                                           :*::**:* ::  :*  **:*:*****:** ::* :**:* :*

M. smegmatis        WP_003897704.1    57  RGTTYLATAPGDLGLARAYVSGDLEPHGVHPGDPYPLLRALAERMEFKRPPARVLANIVR  116
M. marinum          WP_208024746.1    60  RGTTYLATAPGELGLARAYVSGDLQPHGVHPGDPYELLKALTDRVEFKRPSARVLANVVR  119
M. chlorophinolicum WP_048472121.1    60  RGTTYLATAPGDLGLARAYISGNLEAHGVHPGDPYELLNALTEKLDFKRPSARVLAQVIR  119
M. abscessus        WP_005065329.1    57  RGTTYLATAPGDLGLARAYVSGDIEMQGVHPGDPYELLKAMAERLDFKRPPARVLANIVR  116
M. tuberculosis     WP_003899652.1    48  RGATYLATAPGELGLARAYVSGDLQAHGVHPGDPYELLKTLTERVDFKRPSARVLANVVR  107
M. avium            WP_225520609.1    57  RGATYLATAPGELGLARAYVAGDLQAYGVHPGDPYRLLKTLTDRVQFKRPPARVLANVVR  116
M. leprae           WP_041323340.1    60  RGATYLATAPGELGIARAYVAGDLQARGVHPGDPYTLLKTLTDRVQFKRPSLRVLASVVR  119
                                          **:********:**:****::*:::  ******** **.:::::::****  ****.::*

M. smegmatis        WP_003897704.1   117  SIGIEHLKPIAPPPQEALPRWRRIMEGLRHSKTRDAEAIHHHYDVSNTFYEWVLGPSMTY  176
M. marinum          WP_208024746.1   120  SIGVEHLLPIAPPPQETPPRWRRMAEGVLHSKTRDAEAIHHHYDVSNTFYEWVLGPSMTY  179
M. chlorophinolicum WP_048472121.1   120  SIGIEHLKPISPPPQEALPRWRRFAEGLRHSKTRDAEAIHHHYDVSNTFYEWVLGPSMTY  179
M. abscessus        WP_005065329.1   117  SIGLEHLRPIAPPPQEALPRWRRVVEGFRHSKTRDSEAIHHHYDVSNTFYEWVLGPSMTY  176
M. tuberculosis     WP_003899652.1   108  SIGVEHILPIAPPPQEARPRWRRMANGLLHSKTRDAEAIHHHYDVSNNFYEWVLGPSMTY  167
M. avium            WP_225520609.1   117  SIGFERLLPVAPPPQEARPRWRRIADGLMHTKARDAEAIHHHYDVSNRFYELVLGPSMTY  176
M. leprae           WP_041323340.1   120  SLGVEHLLPIAPPPQETPPRWRRITDGLLHSKTRDAEAIRHHYDVSNRFYEWVLGPSMTY  179
                                          *:*.*:: *::*****: *****. :*. *:*:**:***:******* *** ********

M. smegmatis        WP_003897704.1   177  TCACYPTEDATLEEAQDNKYRLVFEKLRLKPGDRLLDVGCGWGGMVRYAARHGVKALGVT  236
M. marinum          WP_208024746.1   180  TCAVYPNEKATLEEAQENKYRLVFEKLRLQPGDRLLDVGCGWGGMVRYAARHGVRVIGAT  239
M. chlorophinolicum WP_048472121.1   180  TCACYQHPEATLEEAQDNKYRLVFEKLRLQPGDRLLDVGCGWGGMVRYAARHGVTAIGVT  239
M. abscessus        WP_005065329.1   177  TCACYPDAEATLEEAQENKYRLVFDKLALKPGDRLLDVGCGWGSMVRYAARRGVRATGVT  236
M. tuberculosis     WP_003899652.1   168  TCAVFPNAEASLEQAQENKYRLIFEKLRLEPGDRLLDVGCGWGGMVRYAARRGVRVIGAT  227
M. avium            WP_225520609.1   177  TCAVYPHPDATLEEAQENKYRLIFDKLRLAPGDRLLDVGCGWGGMVRYAARRGVRAIGAT  236
M. leprae           WP_041323340.1   180  TCAVYPNAEATLEEAQENKYRLIFEKLRLQPGDRLLDVGCGWGGMVRYAARRGVRVIGAT  239
                                          *** :   .*:**:**:*****:*:** * *************.*******:** . *.*

M. smegmatis        WP_003897704.1   237  LSREQATWAQKAIAQEGLTDLAEVRHGDYRDVIESGFDAVSSIGLTEHIGVHNYPAYFNF  296
M. marinum          WP_208024746.1   240  LSAEQATWAQQKISDEGLAQLAEVRHCDYRDVVETGFDAVSSIGMTEHIGVRNYPAYFGF  299
M. chlorophinolicum WP_048472121.1   240  LSKQQAEWAQKAIAEEGLGDLAEVRHSDYRDVRESQFDAVSSIGLTEHIGVANYPAYFRF  299
M. abscessus        WP_005065329.1   237  LSAEQAAWAQKAIADEGLADLAEVRHADYRDTAEREFDAVSSIGLTEHIGIANYPAYFRF  296
M. tuberculosis     WP_003899652.1   228  LSAEQAKWGQKAVEDEGLSDLAQVRHSDYRDVAETGFDAVSSIGLTEHIGVKNYPFYFGF  287
M. avium            WP_225520609.1   237  LSAEQAKWAQRRIDDEGLGDLAQVRHSDYRDVAETGFDAVSSIGLTEHIGVKNYPAYFGF  296
M. leprae           WP_041323340.1   240  LSAEQAKWARKAIDNEGLAEIAQVRYSDYRDIRETEFDAVSSIGLTEHIGVKNYPAYFNF  299
                                          ** :** *.:: : :*** ::*:**: ****  *  ********:*****: *** ** *

M. smegmatis        WP_003897704.1   297  LKSKLRTGGLLLNHCITRPDNRSAPSAGGFIDRYVFPDGELTGSGRIITEAQDVGLEVIH  356
M. marinum          WP_208024746.1   300  LKSKLRTGGLLLNHCITRHDNKSTSFAGGFTDRYVFPDGELTGSGRIITEIQEVGFEVLH  359
M. chlorophinolicum WP_048472121.1   300  LKSKLRTGGLLLNHCITRHDNRHGAAAGGFIDRYVFPDGELTGSGRIITEVQDVGLEVVH  359
M. abscessus        WP_005065329.1   297  LQSRLKTGGLLLNHCITRPDNRTSAVAGYFIDRYVFPDGELTGSGRIITEIQNVGLEVLH  356
M. tuberculosis     WP_003899652.1   288  LKSKLRTGGLLLNHCITRHDNRSTSFAGGFTDRYVFPDGELTGSGRITTEIQQVGLEVLH  347
M. avium            WP_225520609.1   297  LKSKLRTGGLLLNHCITRHDNTSTSFAGGFTDRYVFPDGELTGSGRITCAIQDVGFEVLH  356
M. leprae           WP_041323340.1   300  LKLKLRTGGLLLNHCITRRDNKSTSFAGGFTDRYVFPDGELTGSGRITTEIQEVGLEVLH  359
                                          *: :*:************ **     ** * ****************    *:**:**:*

M. smegmatis        WP_003897704.1   357  EENLRNHYAMTLRDWCRNLVEHWDEAVEEVGLPTAKVWGLYMAGSRLGFETNVVQLHQVL  416
M. marinum          WP_208024746.1   360  EENLRQHYALTLRDWCHNLVEHWDAAVAEVGLPIAKVWGLYMAASRVAFERNNLQLHHVL  419
M. chlorophinolicum WP_048472121.1   360  EENLRNHYAMTLRDWNRNLVEHWDEAVAEVGLATAKVWGLYMAGSRVGFEQNAIQLHQVL  419
M. abscessus        WP_005065329.1   357  EENLRHHYALTLKEWCANLVEHWDEAVAEVGEATAKVWGLYMAGSRLGFERNVVQLHQVL  416
M. tuberculosis     WP_003899652.1   348  EENFRHHYAMTLRDWCGNLVEHWDDAVAEVGLPTAKVWGLYMAASRVAFERNNLQLHHVL  407
M. avium            WP_225520609.1   357  GENFRHHYAMTLRDWCRNLVEHWDAAVDEVGLPTAKVWGLYMAASRVAFEQNNLQLHHVL  416
M. leprae           WP_041323340.1   360  EENFRHHYAMTLRDWCHNLVEHWEDAVAEVGLPIAKVWGLYMAASRVAFEQNNLQLHHIL  419
                                           **:*:***:**::*  ******: ** ***   *********.**:.** * :***::*

M. smegmatis        WP_003897704.1   417  AVKLDDQGKDGGLPLRPWWSA                                        437
M. marinum          WP_208024746.1   420  AANVDTWGE-DXLPLRPWWTA                                        439
M. chlorophinolicum WP_048472121.1   420  AVKLDERGRDGGLPLRPWWTA                                        440
M. abscessus        WP_005065329.1   417  ATKLDERGG-SELPLRPWWQP                                        436
M. tuberculosis     WP_003899652.1   408  ATKVDPRGD-DSLPLRPWWQP                                        427
M. avium            WP_225520609.1   417  AANVDARGD-DDLPLRPWWSP                                        436
M. leprae           WP_041323340.1   420  ATKVDVWGD-NSLPTRPWWTP                                        439
                                          *.::*  *  . ** ****
```

Figure S3

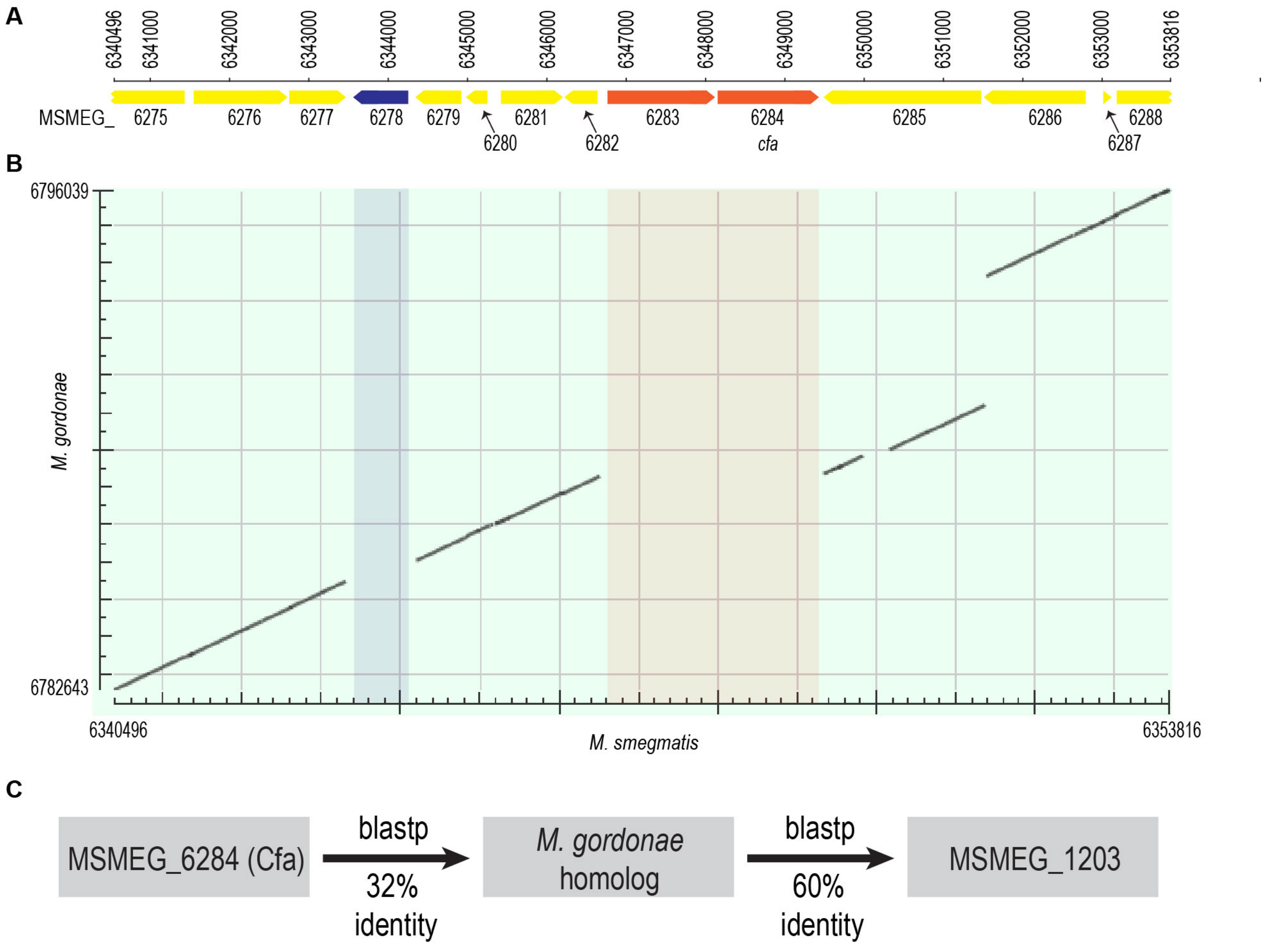
A
6340496
6341000
6342000
6343000
6344000
6345000
6346000
6347000
6348000
6349000
6350000
6351000
6352000
6353000
6353816
MSMEG_
cfa
6285
6286
6287
6288
B
6796039
6782643
M. gordonae
6340496
M. smegmatis
6353816
C
MSMEG_6284 (Cfa)
blastp
32%
identity
M. gordonae
homolog
blastp
60%
identity
MSMEG_1203

Figure S4

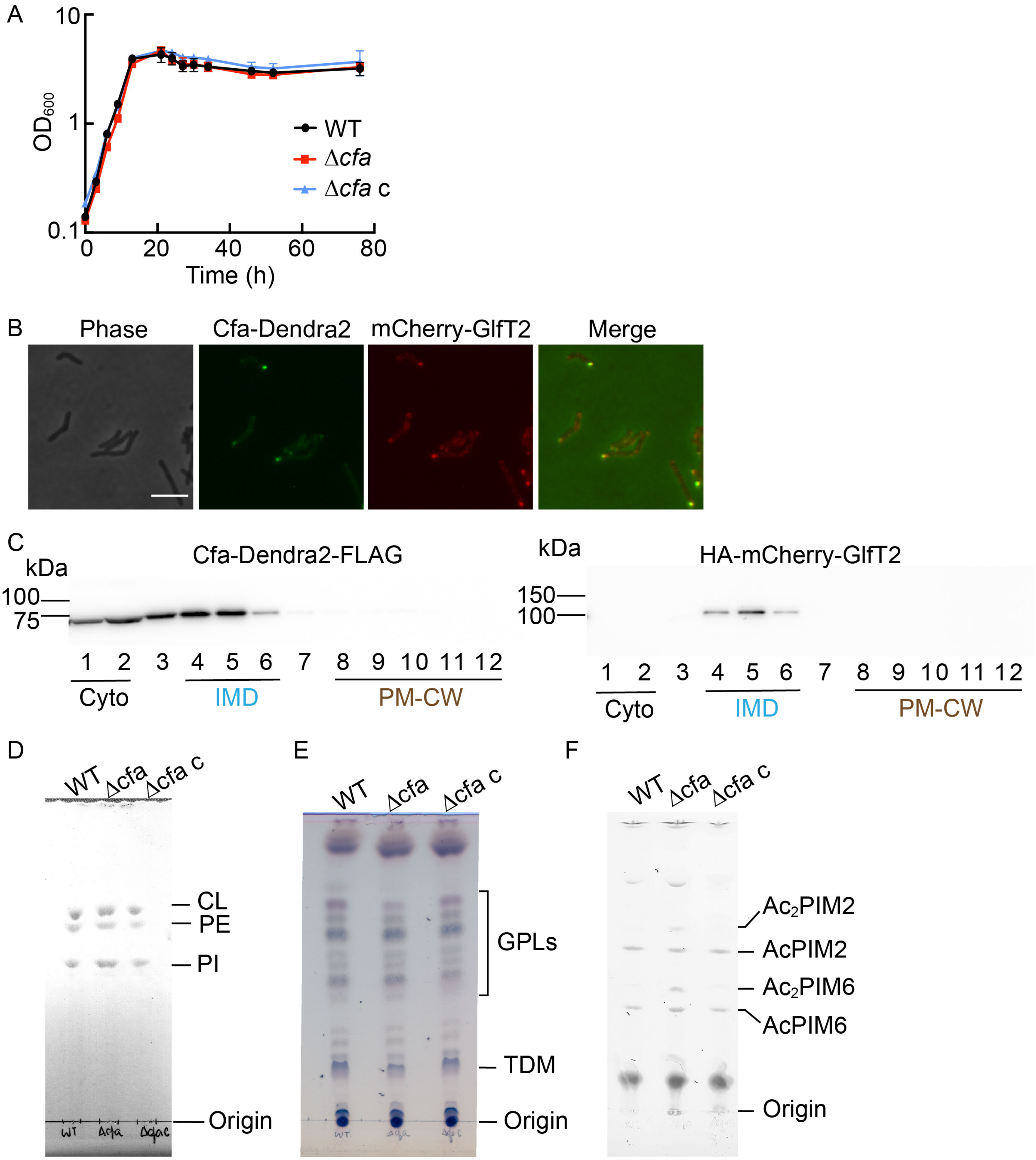
A
10
1
0.1
OD600
0
20
40
60
80
Time (h)
WT
Δcfa
Δcfa c
B
Phase
Cfa-Dendra2
mCherry-GlfT2
Merge
C
kDa
Cfa-Dendra2-FLAG
100
75
1 2 3 4 5 6 7 8 9 10 11 12
Cyto
IMD
PM-CW
kDa
HA-mCherry-GlfT2
150
100
1 2 3 4 5 6 7 8 9 10 11 12
Cyto
IMD
PM-CW
D
WT
Δcfa
Δcfa c
CL
PE
PI
Origin
E
WT
Δcfa
Δcfa c
GPLs
TDM
Origin
F
WT
Δcfa
Δcfa c
Ac2PIM2
AcPIM2
Ac2PIM6
AcPIM6
Origin

Figure S5

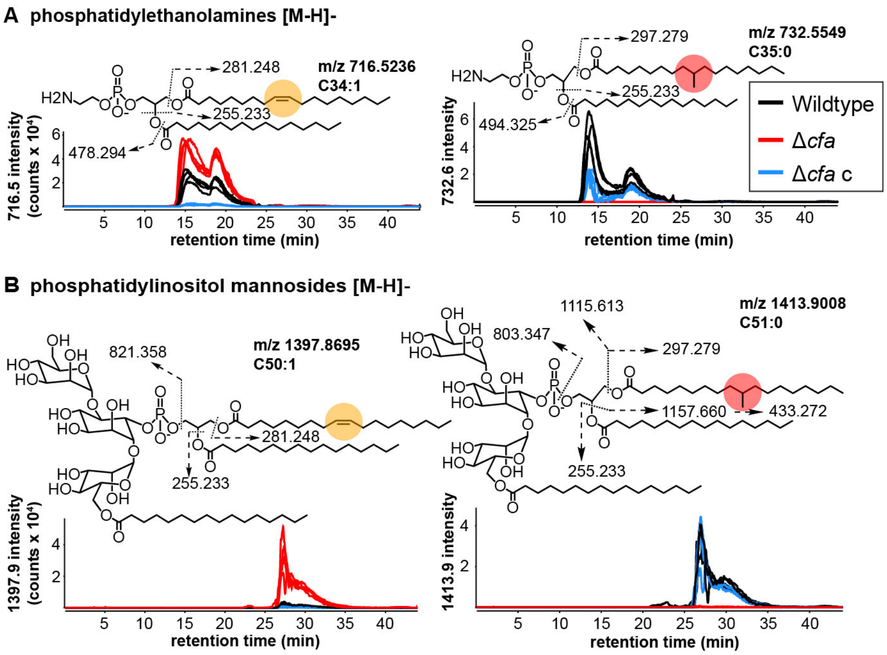

A phosphatidylethanolamines [M-H]-
281.248
m/z 716.5236
C34:1
H2N
255.233
478.294
716.5 intensity (counts x 10⁴)
retention time (min)
297.279
m/z 732.5549
C35:0
255.233
494.325
732.6 intensity
Wildtype
Δcfa
Δcfa c
B phosphatidylinositol mannosides [M-H]-
821.358
m/z 1397.8695
C50:1
281.248
255.233
1397.9 intensity (counts x 10⁴)
1115.613
m/z 1413.9008
C51:0
803.347
297.279
1157.660
433.272
255.233
1413.9 intensity

### Figure S6

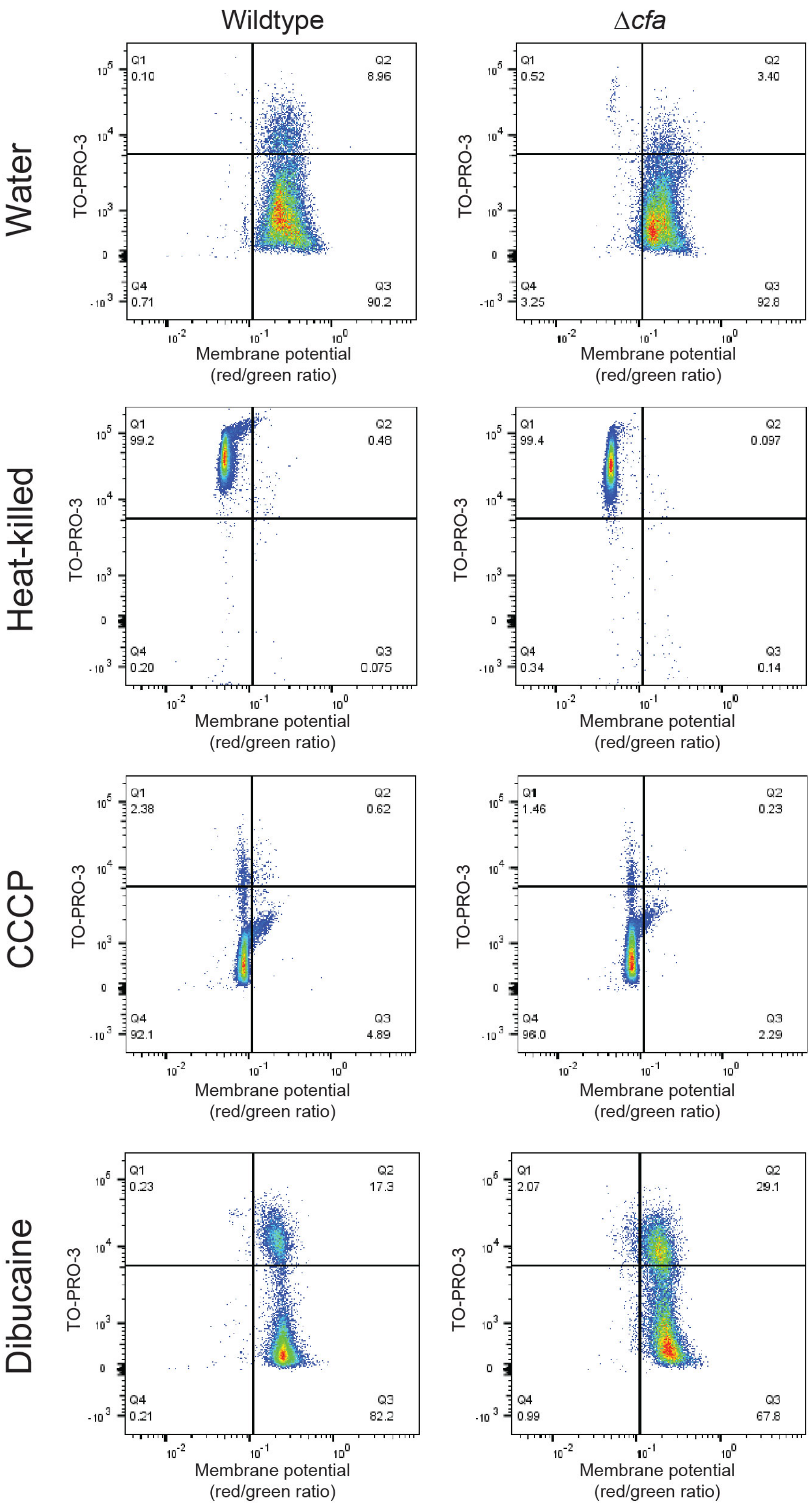

Wildtype
Δcfa
Water
Heat-killed
CCCP
Dibucaine
TO-PRO-3
Membrane potential (red/green ratio)
Q1 0.10
Q2 8.96
Q4 0.71
Q3 90.2
Q1 0.52
Q2 3.40
Q4 3.25
Q3 92.8
Q1 99.2
Q2 0.48
Q4 0.20
Q3 0.075
Q1 99.4
Q2 0.097
Q4 0.34
Q3 0.14
Q1 2.38
Q2 0.62
Q4 92.1
Q3 4.89
Q1 1.46
Q2 0.23
Q4 96.0
Q3 2.29
Q1 0.23
Q2 17.3
Q4 0.21
Q3 82.2
Q1 2.07
Q2 29.1
Q4 0.99
Q3 67.8
